# Sex-specific chemosensory gene expression in the whitelined sphinx moth (Lepidoptera: *Hyles lineata*) suggests a role for odorant-binding proteins in host plant choice

**DOI:** 10.1101/2024.09.04.611314

**Authors:** R. Keating Godfrey, Anthony Auletta, Edison Cheung, Riley Harper, Kireina Kates, Akito Y. Kawahara, Yichen Li, Cristina Mercado, Fernando Miguelena, Ginger Pickett, Peter DiGennaro

## Abstract

The whitelined sphinx moth, *Hyles lineata*, is a generalist during both the larval and adult stages with a broad geographic range extending across North and Central America. Within the genus *Hyles* there have been multiple transitions to a narrower host plant range, making *Hyles* an ideal group to study the evolution and mechanisms of host plant selection. We characterize sex- and appendage-specific chemosensory gene expression in *H. lineata*, the oldest extant member of the genus. We also describe female-specific gene expression in appendages used to sense plant surfaces as a means of identifying candidate genes involved in host plant choice. Sensilla on these appendages house sensory neurons and support cells that express chemosensory genes, receptors, and small proteins that bind, shuttle, and transport small molecules to allow detection of odorants and other small molecules. We considered genes detected more frequently in the female leg and ovipositor samples to be candidate oviposition-relevant genes. Most chemosensory genes of interest were detected in both sexes, while several odorant receptors were only detected in females. We identified 18 putative chemosensory genes that were specific to female legs, ovipositors, or both body parts. However, most of these genes did not reach statistical criteria to be considered differentially expressed. Instead, a set of OBPs show statistically significant female-biased expression in legs and ovipositors. These genes may serve as candidates for future study of the evolution and mechanisms of oviposition behavior in this species and its relatives.

**Summary:** The sphinx moth genus *Hyles* contains both generalist and specialized feeders, making it an ideal system to study the evolution of host plant breadth. The whitelined sphinx moth (*Hyles lineata*) is a generalist feeder at both adult and larval stages and is the oldest lineage of the genus. In this analysis of sex-specific gene expression, Godfrey and colleagues identify several odorant binding proteins as being more frequently detected in female appendages used to assess host plants. These genes could play a role in host plant selection and can be targets for future mechanistic studies on this species.

## Background

Phytophagous insects comprise at least a half million species, nearly 25% of described multicellular animals (1), and the interactions among related insects and their host plants are thought to drive trait evolution in both groups (2–4). Studies of broad patterns of adaptation and coevolution provide a strong theoretical framework for characterizing the driving forces in insect-host plant associations (5,6). Simultaneously, detailed study of insect behavior indicates not only inter-but also intraspecific variation in insect associations with host plants (7) Specialists that feed on one or a few plant genera show the greatest consistency in host plant association, but even in these cases, individuals may vary in preference (8). These complex interactions result from genetic and geographic variation in the distributions of insects and their host plants interacting with developmental and ecological variables. These systems provide exciting opportunities to understand how local dynamics drive natural selection and affect patterns of evolution.

While the stage is set to understand broad patterns of adaptation and coevolution in plant-insect interactions, characterizing specific interactions is an important component of bringing these patterns into focus. One of the first steps in any insect-host plant association is a female ovipositing, or laying eggs, onto or into plant tissue (9). She decides whether to oviposit using sensory receptors on her appendages to assess host plant qualities including surface texture and chemical composition. Butterflies and moths (Lepidoptera) use a combination of olfactory and visual cues to locate and identify host plants from a distance (10–13), and rely on mechanosensation and gustation — or a combination of these senses — to make the final decision about whether or not to lay eggs (13–16). Moth proboscises, legs, and genitalia are known to serve sex-specific contact chemosensory functions (e.g., 15,17,18) Sensilla on these appendages house sensory neurons and support cells that express chemosensory genes, receptors and small proteins that bind, shuttle, and transport small molecules to allow detection of odorants and other small molecules. Moths have been shown to use sensory receptors on their antennae (19), tarsi of their fore- and mid-legs (20), and ovipositor (15,21) to contact and assess the plant surface.

The penultimate behavior of laying an egg is probing the plant surface with the ovipositor (10). Moth ovipositors are covered in setae, some of which are sensilla that express olfactory receptors (ORs), gustatory receptors (GRs), or ionotropic receptors (IRs) (15,16,22,23). These chemosensory receptors, along with sensory neuron membrane proteins (SNMPs) and odorant-binding proteins (OBPs), are core components of chemosensory transduction in insects. Females assess host plant suitability and quality using volatile and nonvolatile compounds secreted from plant tissue, and plant compounds that act as oviposition stimulants underlie within-species differences in host plant preference (19). Contact of the ovipositor with the plant surface provides an insect the opportunity to assess compounds such as salts, sugars, and plant secondary metabolites that are not readily detected at a distance. Volatile compounds can also be sensed in close contact with the plant; therefore, olfaction and gustation are likely used in the decision to lay an egg.

Insect odorant receptors evolved from gustatory receptors (24,25) and, while they are characterized as responding to volatile and nonvolatile compounds, respectively, this distinction is not always so clear and functional overlap exist (reviewed in 26). Additionally, several IRs, which are not related to GRs but instead to ionotropic glutamate receptors (iGluRs), are multimodal and respond to temperature, tastants, and odorants (Wicher and Miazzi, 2021). OBPs are expressed in olfactory and gustatory sensilla and bind molecules to help solubilize them into the hemolymph.

Additionally, they transport molecules and buffer against rapid changes in odorant levels (reviewed in 27). SNMPs are transmembrane proteins involved in the recognition and transport of hydrophobic molecules. There are at least three subfamilies of SNMPs in insects (28), with SNMP1 implicated in pheromone detection in Lepidoptera (29,30).

Here we focus on a hawk moth, the whitelined sphinx (Sphingidae*: Hyles lineata*) to study oviposition-relevant chemosensory genes. Hawk moths are large-bodied, fast-flying moths that are known both as defoliating agricultural pests (31,32) and important pollinators of night-flowering plants (33–35). Additionally, there are well-supported phylogenies at the family level (36) and for several clades within Sphingidae (37–39), making this an opportune group to link genotypic evolution with phenotype. *Hyles* is a particularly group to study host-plant associations because it is the earliest diverging lineage and a broad host-plant generalist. Within more recent lineages of the genus, multiple transitions to host specialization have occurred, including at least two transitions to plants containing toxic secondary compounds (37).

In this study, we quantified gene expression in *H. lineata* sensory appendages in order to generate a list of candidate oviposition-relevant chemosensory genes for future experiments. We focused on the legs and ovipositor, which are thought to make direct contact with the leaf surface just before oviposition. The sensory machinery used for oviposition likely overlaps with that used for behaviors not specific to females, but genes expressed only in females or more frequently in females are likely important to female-specific behaviors. Gene expression was therefore compared across males and females to determine female-biased expression in the legs and genitalia. Genes that are leg- or ovipositor-specific and detected more frequently in females are considered good candidates for ones used in oviposition behavior. We additionally characterized gene expression in the proboscis because moths use this appendage to probe a another part of the plant, the inflorescence, for nectar. This is not a sex-specific behavior, as both males and females nectar, so we expected to find fewer differences in chemosensory genes between sexes than for other body parts.

## Methods

### Specimen collection

*Hyles lineata* were collected from Pima County, Arizona in April and September, 2022. Specimens were collected by attracting moths to two 50 watt mercury vapor bulbs or a 50 watt UV LED panel light were directed at a reflective, white sheet from sunset until ∼23:00. Moths that landed on the sheet were sexed and stored individually in 50 mL conical falcon tubes in a cool, dark location until being placed directly in a -80 °C freezer. Moths were preserved this way within 4 hours of collection. Four individuals of each sex were used for gene expression experiments. Collection dates and locations are available in Table S1.

### RNA Isolation

RNA was isolated from the proboscis, the legs (tibia and tarsal segments), and genitalia, using a TRIzol-chloroform isolation procedure. For sample information see Table S2. For some specimens fewer than six legs were recovered, but at least one fore-, middle-, and hind leg was included for each specimen. First, a moth was removed from the -80 °C freezer and placed in a petri dish on ice. Each of the three body parts were excised using microscissors and placed directly via forceps in a 1.5 mL RNAse-free microcentrifuge tube on ice. Each tube contained 800 µL of cold TRIzol (Thermofisher cat. # 15596026) and two autoclaved stainless steel homogenizer beads (4 mm). For females, only genitalia segments that included the ovipositor were excised for processing, but we use the term “genitalia” for the comparison of sexes. Samples were homogenized using a bead beater cooled to -20 °C for 2.5 minutes at 1900 RPM. Following homogenization, 200 µL of ice-cold chloroform was added to each sample, mixed by inversion, and incubated at room temperature for 3 minutes. Samples were centrifuged for 15 minutes at 12,000 x g and 4 °C, and the upper phase of the sample was transferred to a new tube. RNA was precipitated with ice-cold 100% isopropanol and 1 µL mussel glycogen (Milipore cat. # 361507) and then washed twice with room temperature 70% ethanol. The resulting pellet was dried before being resuspended in 50 µL of RNAse-free water and stored at -80 °C until sequencing.

### Sequencing

RNA quality control, library preparation, and sequencing were performed by the University of Florida’s Interdisciplinary Center for Biotechnology Research’s Gene Expression and Genotyping core facility (ICBR; RRID:SCR_019145) using a 2 x 150 paired-end sequencing format on a NovaSeq6000 (Illumina; San Diego, CA, USA).

### Read Quality Assessment and Differential Gene Expression Analysis

Preliminary data analysis for this project was conducted by 31 students in the University of Florida’s Entomology Department’s course-based undergraduate research experience (CURE): Insect Research and Scientific Engagement. Students worked in teams to test hypotheses about body part and sex-specific expression, using pipelines that were later refined for final data analysis and visualization.

Reads were trimmed using Trimmomatic v.0.39 (40; RRID:SCR_011848) and read quality was assessed by inspection of FastQC v.0.12.1 results (RRID:SCR_014583). Reads were mapped to the indexed genome using HISAT2 v.2.2.1.3n (41; RRID:SCR_015530). Following read quality assessment, we calculated counts for each gene using a published, structural annotation of the *H. lineata* genome (42, see details in Chemosensory Gene Annotation) and htseq-count function of HTSeq v2.0.3 (43; RRID:SCR_011867). This structural annotation includes multiple transcripts for a number of genes, but we limited counts to primary transcripts. We assessed whether a gene was likely present in a sample by calculating transcripts per million (TPM) for each gene (presence/absence analysis). We then performed differential gene expression (DGE) across samples by sex and body part using DESeq2 v3.18 (44; RRID:SCR_015687) in R (v. 4.3.3 (2024-02-29) -- "Angel Food Cake") with RStudio (v. 2023.12.1+402 "Ocean Storm") and visualization using ggplot2 (45). We removed low counts from our dataset by limiting analysis to genes that had at least 30 counts in three or more samples. We grouped the body part and sex variables to produce the generalized linear model y ∼ group, which allowed us to compare each sex + body part combination with any other. To identify chemosensory genes potentially involved in egg-laying behaviors, we first compared gene expression across body parts of each sex, with a particular interest in genes recovered more frequently from female legs and genitalia. To further assess ovipositor-specific expression in females, we compared gene expression between female genitalia and either female legs or proboscises, respectively.

### Chemosensory Gene Annotation

To determine if there are sex-specific differences in chemosensory gene expression, we sought to first identify genes important to chemosensory behavior from the *Hyles lineata* genome. We previously performed a structural annotation of the *H. lineata* genome (42) using BRAKER2 (46; RRID:SCR_018964) with RNA-seq evidence from a caterpillar and two adult samples, one of each sex. We identified putative ORs, GRs, and IRs, OBPs, and SNMPs from the existing structural annotation. We used two approaches to determine which transcripts might be chemosensory genes. First, we used the functional annotation tool eggNOG-mapper (47,48; RRID:SCR_021165), which uses orthologous groups (OGs) to determine the putative function of genes. We filtered results for genes that contained the conserved, seven-transmembrane domains of insect olfactory receptors and gene ontology (GO) terms for chemosensation, which includes those specific to olfaction and gustation. Second, we blasted NCBI RefSeq protein sequences from a related hawk moth, the tobacco hornworm (*Manduca sexta*), and the well-characterized fruit fly (*Drosophila melanogaster*), against proteins predicted from the *H. lineata* genome annotation. We then combined lists of putative chemosensory genes from the eggNOG mapper annotation and blast results (N = 106) and used the NCBI BLASTP tool (RRID:SCR_001010) to search NCBI’s clustered, non-redundant (nr) protein database for matches with characterized function. This is a database formed by gene clusters showing sequences within 90% length and 90% identity to other sequences of the cluster. Sequences from our list that returned a chemosensory gene cluster were considered likely chemosensory genes for differential gene expression analyses (N = 85, Table S3).

Following differential gene expression analysis, we constructed gene trees of putative odorant-binding protein genes that were recovered more frequently from the female legs and ovipositor samples. We used AliView to align *H. lineata* sequences with OBP sequences from the NCBI non-redundant protein database for the sphingid moth *M. sexta*, the related bombycoid silk moth, *Bombxy mori*, and *D. melanogaster*. We built phylogenies using the ape (49) and phangorn (50) packages in R. First, we used the *dist.ml* function with the WAG model, which uses a maximum-likelihood approach informed by 182 protein families to model amino acid substitution (51), to generate a distance matrix between sequences. We then built an unrooted tree using the neighbor-joining (*nj*) function from ape and visualized it using ggtree and msaplot (52–54).

## Results

### Sample Quality

Sequencing resulted in an average of 44.9 million reads per sample (sd = 11, N = 25; Table S2). The majority of sample reads mapped to the *H. lineata* genome at rates of > 82%; however, one female proboscis sample (sample ID Hl22042008) showed mapping < 55% (Table S2), and this sample was excluded from downstream analyses. While results from blastn against the entire NCBI database show the most hits for the small elephant hawk moth (*Deilephila porcellus*), the second most frequent matches are from common ivy (*Hedera helix*). Thus, given that pollen is deposited on the proboscis during foraging, it is possible that the low mapping percent of this sample is from plant contamination. Otherwise, our samples show high levels of pairwise correlation and cluster in gene expression space according first to sex and then to body part, with the same body parts from each sex closer in gene space (Figure S1).

### Chemosensory Gene Annotation

The eggNOG-mapper tool identified orthologs for 16,222 of the 20,268 genes predicted from the structural annotation. There were 14,489 primary transcript genes in this set and we proceeded with these for chemosensory annotation. BRAKER2 annotates multiple transcripts for the same gene in some cases, but eggNOG-mapper identified the same ortholog for all transcripts where multiple were present, so limiting our analysis only to primary transcripts should not reduce our ability to identify chemosensory genes. We recovered 2 SNMP, 11 OBP, 13 IR, 11 GR, and 48 OR sequences from a structural annotation of the *H. lineata* genome (Figure S2A). These are likely conservative estimates of total members in each gene family in this species but provided enough information to assess differences in well-identified, highly expressed chemosensory proteins. Our functional annotation recovered two copies of a well-characterized, single-copy ortholog, the odorant receptor co-receptor (*orco*; 55). This led to further inspection of the genome and we found that these *orco* sequences are located on two contigs that show nearly perfect synteny along 1250 kb of their lengths (Figure S2B), suggesting this is a region that was not properly collapsed during assembly. Importantly, we recovered both sequences from all samples in our dataset, indicating o*rco* is present in all sensory structures sampled. *Sex and body part-specific differences in gene expression*

When we looked at genes expressed in ≥ 3 samples of each body part of each sex, we found 47 chemosensory genes expressed in the genitalia (19 shared, 28 female-specific; Figure 1A), 38 expressed in the legs (21 shared, 10 female-specific, 7 male-specific; Figure 1B), and 32 expressed in the proboscis (15 shared, 16 female-specific, and one male-specific; Figure 1C). In comparisons across female and male genitalia and legs, 18 genes were exclusively detected in the ovipositor and female legs, 12 of which were ovipositor-specific (Figure 1D; for identification of these genes see Identification of oviposition-relevant chemosensory genes section below).

**Figure 1.**
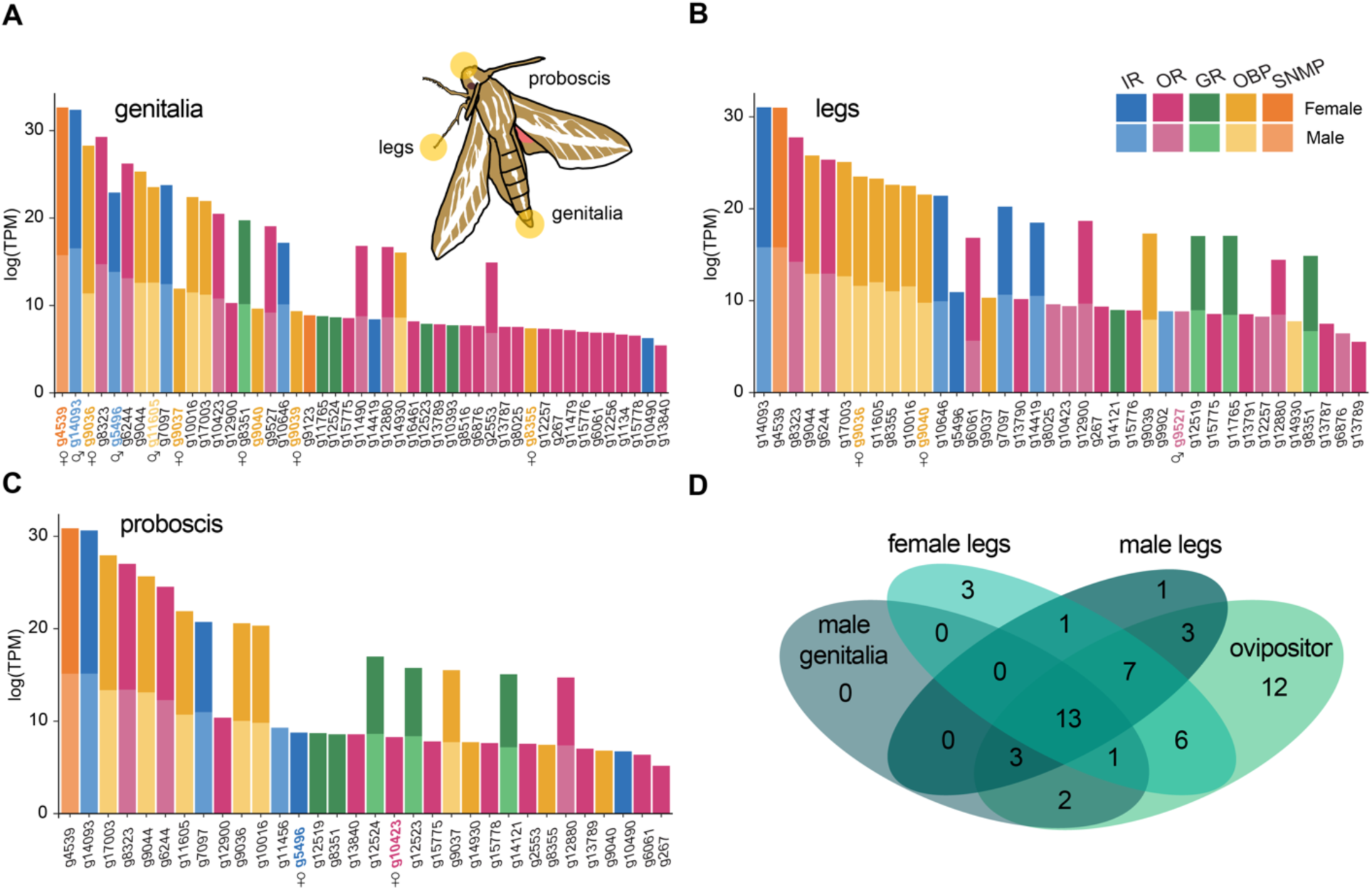
Detection of chemosensory genes in male and female H. lineata genitalia (A), legs (B), and proboscis (C). Gene counts in transcripts per million (TPM) for genes detected in ≥ three samples of each sex and body part, ranked by female expression level. Gene IDs from structural annotation listed along the x-axis. Where present, sex symbols below IDs indicate genes detected more frequently in the respective sex in DGE analysis. (D) Venn diagram displaying the number of genes unique to female legs and ovipositor. GR = gustatory receptor, IR = ionotropic receptor, OR = olfactory receptor, OBP = odorant binding protein, SNMP = sensory neuron membrane protein.

Filtering data for low counts resulted in 12,621 genes for DGE analysis (Table S4). DESeq2 removed another 3,070 genes either because they had low mean normalized counts or were extreme outliers (Love et al., 2014). While most GRs, IRs, OBPs, and SNMPs were recovered from both sexes, in our presence or absence analysis using TPM, nearly one-quarter of OR genes were detected only in female samples (Figure S2A). In the full DGE model we did not observe any obvious patterns in chemosensory gene expression by sex or body part (Figure S3), but in pairwise comparisons between male and female body parts OBPs frequently showed female-biased expression (Figure 2). Five OBPs show greater expression in female ovipositors when compared with male genitalia: *g9036*, *g9037*, *g9040*, *g9039*, and *g8355*. Two of these, *g9036* and *g9040*, also show greater expression in female legs. We recovered a putative ortholog of SNMP1, *g9123*, from all tissues except the male proboscis and did not detect DE across body parts. However, an ortholog of SNMP2, *g4539*, showed significantly greater expression in the *H. lineata* ovipositor as compared with male genitalia. In proboscis comparisons, one IR, *g5496*, and one OR, *g10423*, were detected more frequently from females, and we did not detect any DE chemosensory genes in male proboscises; notably, this tissue had a lower number of significant DEGs than other comparisons (Figure 2).

**Figure 2.**
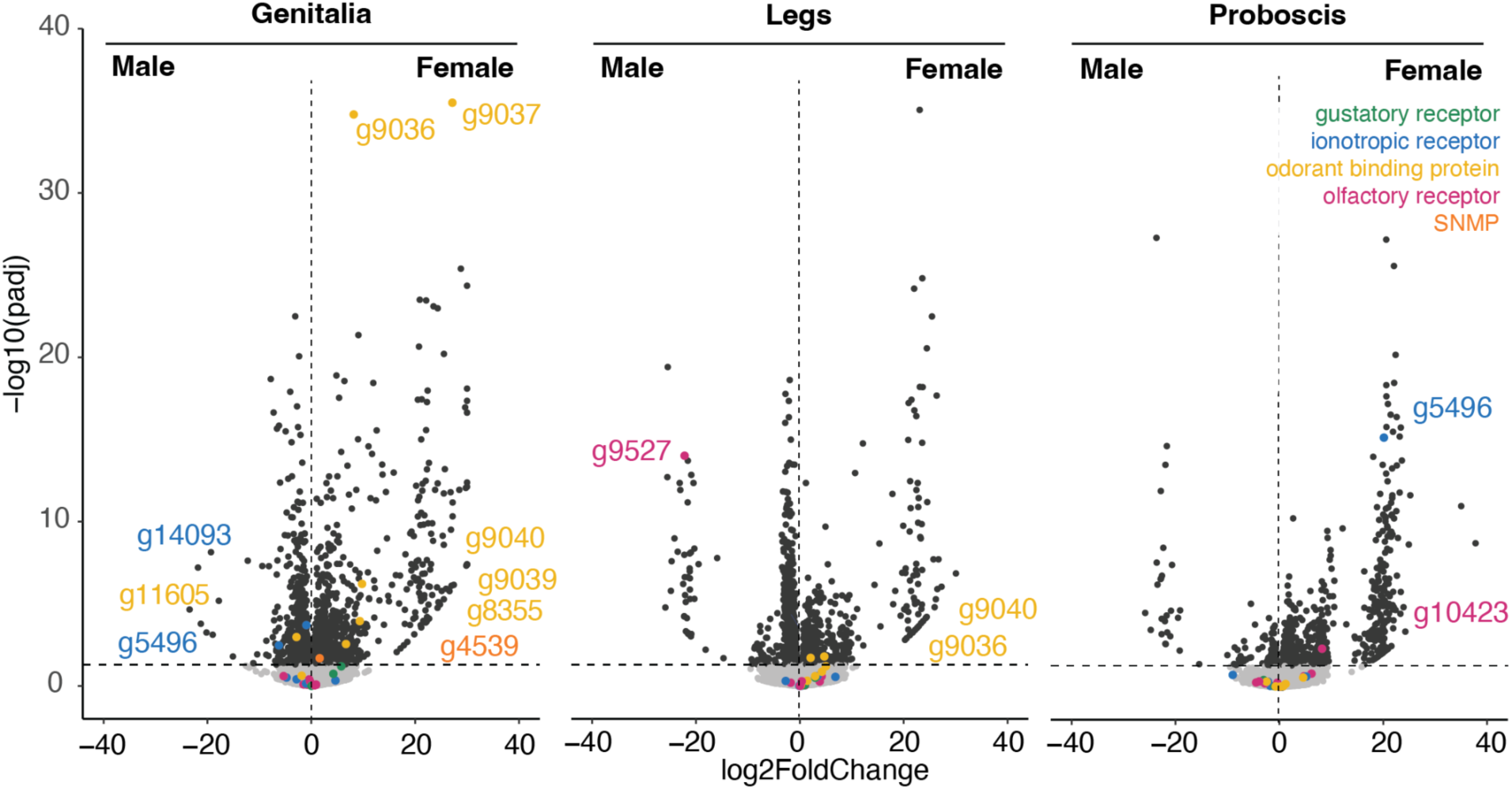
Differential gene expression between male and female body parts. Chemosensory genes of interest indicated by colors and labels, and other significant genes indicated in dark gray. The horizontal dotted line indicates an adjusted p-value of < 0.05. The vertical dotted line placed where log2FoldChange = 0.

### Identification of oviposition-relevant chemosensory genes

Although a detailed ethogram of *H. lineata* oviposition behavior is not yet reported, based on published behavioral observations of the hawk moth *M. sexta* (20) and personal observations from lab colonies, it is likely that females use their legs and ovipositor to assess surfaces prior to laying eggs. Therefore, we considered chemosensory genes present in these body parts as potentially relevant to oviposition behavior. When we compared genes in these sensory appendages across sexes, we found 12 chemosensory genes specific to ovipositor samples: seven putative ORs (*g1134*, *g8516*, *g15778*, *g11479*, *g12256*, *g13840*, *g16461*), three putative GRs (*g10393*, *g12523*, *g12524*), one SNMP (*g4539*), and one IR (*g10490*; Figure 1C). Six additional genes were detected only in female legs and ovipositors: seven ORs (*g267*, *g15775*, *g15776*, *g13787*, *g13789*) and one OBP (*g9037*). Only one of these female-specific genes, OBP *g9037*, was statistically significant in DGE analysis (Figure 2).

When we looked for ovipositor-specific expression in females, we found that *g9036* and *g9037*, two OBPs identified in male versus female DGE comparisons, showed overrepresentation in the ovipositors when compared with the proboscis or legs (Figure 3). One OR, *g9527* was also detected more frequently in the ovipositor than in the legs. A separate set of OBPs, including *g8355*, *g9040*, and *g11605*, showed greater expression in the legs when compared with the ovipositors. The leg and proboscis samples both showed greater expression of the putative OBP *g17003* and IR *g14093* than the ovipositor.

**Figure 3.**
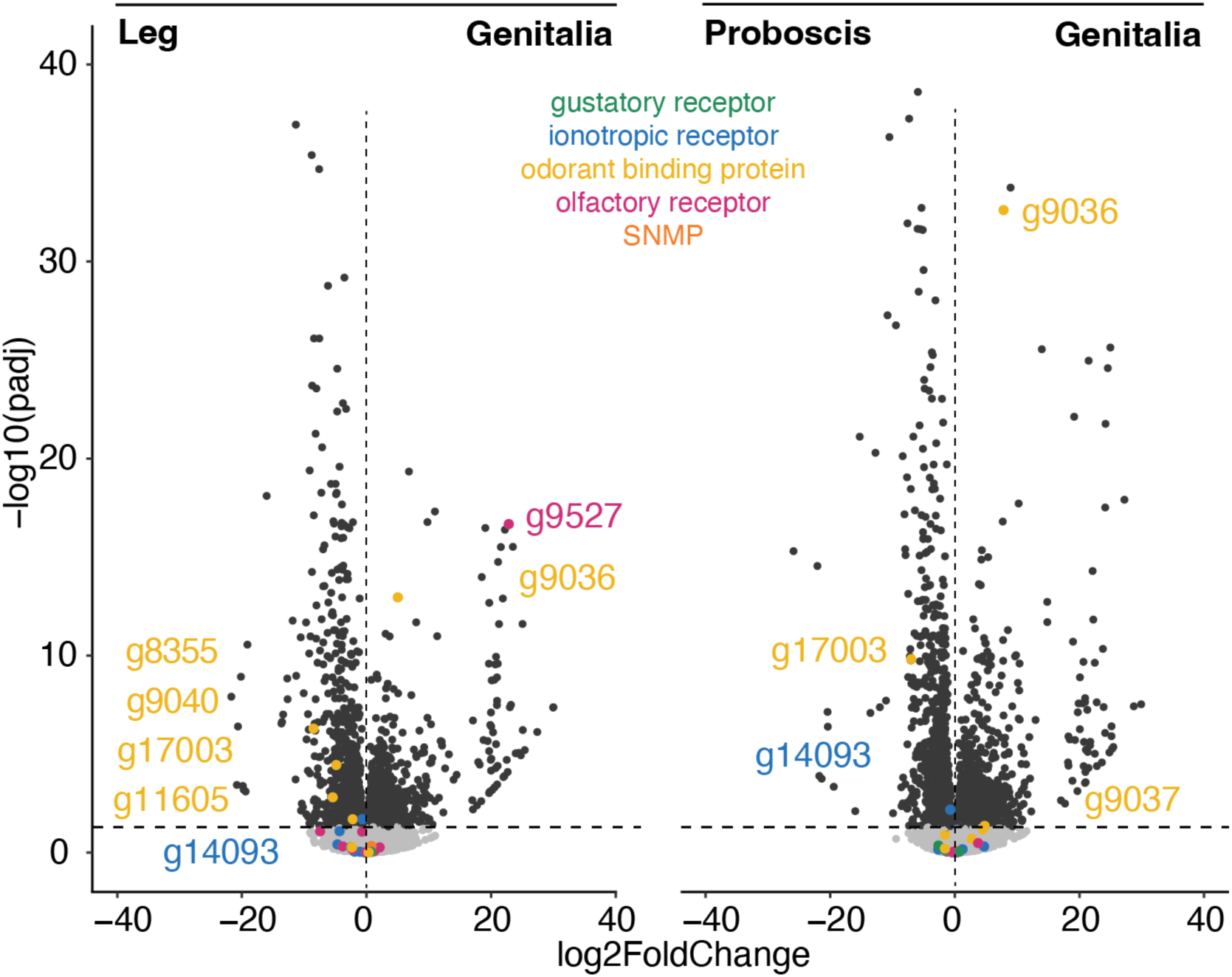
Differential gene expression between female body parts. Odorant binding proteins and one olfactory receptor recovered more frequently from female genitalia than other female sensory structures. Significant chemosensory genes are indicated by the colors and labels. The horizontal dotted line indicates an adjusted p-value of < 0.05. The vertical dotted line indicates that the log2FoldChange = 0.

### Odorant-binding proteins

To better understand the identities of *H. lineata* OBPs, a number of which showed greater expression in females, we assessed sequence similarity with OBPs characterized from two related bombycoid moths, *M. sexta* (*Msex*) and *Bombxy mori (Bmor)*, along with *D. melanogaster* (*Dmel*; Figure 4). Three *H. lineata* genes, *g9037*, *g9039*, and *g9040* fall into a clade that includes orthologs of one of the better-characterized OBPs, *LUSH* (27,56,57). The *g14930* OBP gene, which showed high levels of expression in all sensory appendages (Figure 1A, 1B, 1C), and significantly higher expression in male legs (Figure 2B), is closely related to *DmelOBP83a* and *DmelOBP83b*, which are expressed in antennal chemosensory sensilla in the fly (27,58). Two OBPs, *DmelOBP73a* and *DmelOBP59a*, are highly conserved OBPs with orthologs in most insect taxa (59,60). Here we identified *g10016* as being related to *DmelOBP73a*, but did not recover any OBPs as falling into the *DmelOBP59a* clade (Figure 4). Additionally, *H. lineata* genes *g9044* and *g9036* appear to fall within a moth-specific clade, but this contains *MsexGOBP19d* and *MsexGOBP28a*, which are orthologs of *D. melanogaster* OBPs.

**Figure 4.**
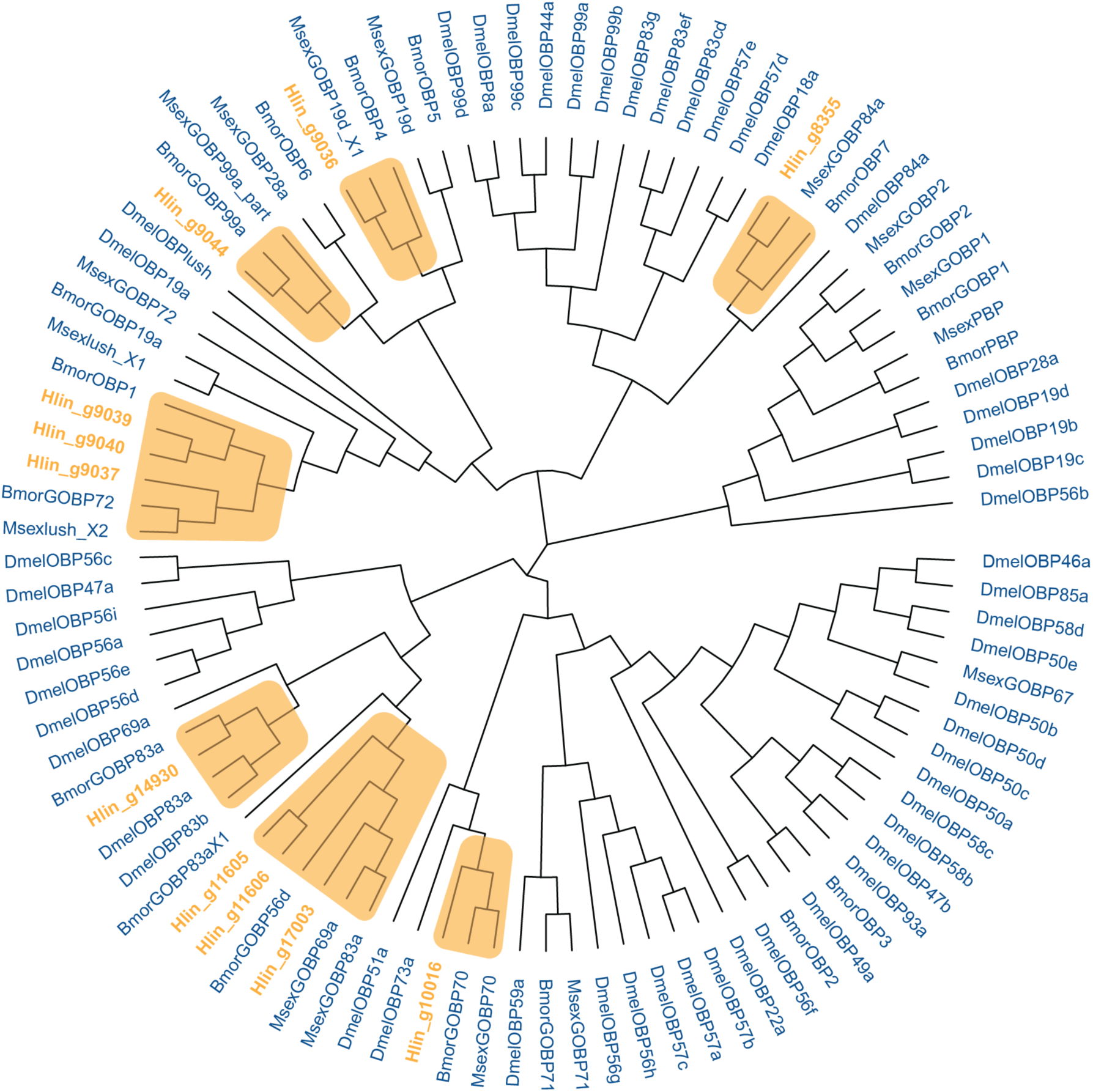
Sequence similarity of *H. lineata* putative odorant-binding proteins in an unrooted, neighbor-joining tree with sequences from *M. sexta*, *B. mori*, and *D. melanogaster*. *H. lineata* sequences are bolded and orange in color, with shaded boxes indicating clades they fall within. DmelOBP = *D. melanogaster* odorant-binding protein, BmorOBP = *B. mori* odorant-binding protein, Msex = *M. sexta* odorant-binding protein.

## Discussion

In this study, we assessed chemosensory gene expression in appendages of both sexes of the whitelined sphinx moth to identify oviposition-relevant genes. We considered genes more frequently detected in female leg and ovipositor samples to be candidate, oviposition-relevant genes. Most GRs, IRs, OBPs, and SNMPs were detected in both sexes, while a number of ORs were only detected in females. Since genes expressed in the mouthparts are likely more involved in nectaring than host plant selection, we contrast patterns of chemosensory gene expression in the mouthparts and in males with those expressed in the female legs and ovipositors. We identified 18 putative chemosensory genes that were specific to female legs, ovipositors, or both body parts (not detected in female mouthparts or male samples). However, most of these genes did not reach statistical criteria to be considered differentially expressed. Instead, a set of OBPs showed statistically significant female-biased expression in legs and ovipositors. These genes may serve as candidates for future study of the evolution and mechanisms of oviposition behavior in this species and its relatives.

### Chemosensory genes

Overall, we recovered fewer chemosensory genes for each protein type of interest (except SNMPs) than has been described from other moth genomes. For example, the related hawk moth *M. sexta*, has 2 SNMPs, 18 OBPs, 21 IRs, 45 GRs, and 75 ORs (61), while the silk moth *B. mori* has 2 SNMPs, 45 OBPs (62), 76 GRs (63), and 66 ORs (64). Chemosensory genes evolve rapidly and show lineage-specific expansions in insects (e.g., 25,65), so the differences in our estimates for *H. lineata* could represent receptor evolution. However, we suspect our numbers indicate that our functional annotation is conservative. This is supported by the fact that we recovered only one of two highly conserved insect OBP orthologs. Thus, we have high confidence in the genes we recovered as showing sex-specific expression, but a more detailed annotation could reveal additional differentially expressed chemosensory genes.

### Body-part and sex-specific chemosensory gene expression

Sex-specific behaviors are thought to be an important factor in sensory gene expression differences between males and females (e.g., 17), with females likely expressing sensory genes specific to host plant selection and egg-laying behaviors. Conversely, while there are subtle sex differences in foraging, males and females feed from the same set of nectar sources (66), and we expected there to be a great deal of overlap in genes expressed in the proboscises. Indeed, these samples showed the greatest parity between sexes, with no female- or male-specific GRs and very few other sex-specific chemosensory genes detected. In fact, the proboscis is the only tissue from which we recovered the exact same set of GRs from both sexes (Figure S2). Because of its role in feeding, we expected a number of GRs might show proboscis-specific expression, but instead found broad overlap in GRs across tissue types.

Insect tarsi and genitalia are covered in sensilla, many of which are used for airborne and contact chemoreception (67–70). While the sequence of oviposition choice behaviors is not well-characterized in *H. lineata*, other moths and butterflies are known to use dragging or drumming behavior of these appendages on the plant surface before deciding whether or not to oviposit (71,72). For moths, contact chemoreception seems particularly important in the final decision to oviposit (73). Therefore, we expected to identify female-specific GR and OR expression in the tarsi and ovipositor. Two putative GR genes, *g12518* and *g12523*, were detected only from female leg and ovipositor samples, but neither were differentially expressed. These genes show the greatest sequence similarity to the *M. sexta* gustatory receptor 64f and are located in a cluster of at least five putative GRs in the *H. lineata* genome (*g12518*, *g12519*, *g12521*, *g12523*, and *g12524*).

Expression of the ortholog DmelGR64f in legs of *D. melanogaster* is required for sucrose preference (74), and these GR orthologs are likely sugar receptors. An additional eight putative ORs were detected only from these tissues, but also were not differential expressed. Our inability to detect differences may be due to low sample sizes. Repeated experiments will help to more confidently determine whether these receptors are candidates for future mechanistic and evolutionary studies.

In addition to receptors, the sensation of plant compounds requires proteins that bind and transport ligands such as sensory neuron membrane proteins and odorant binding proteins. It is in these classes of genes that we detected higher expression in females, suggesting these might be important for oviposition decisions. Specifically, we detected SNMP2 and five OBPs as occurring more frequently in the ovipositor, with two of these OBPs also showing greater expression in female legs. SNMPs likely play a role in pheromone sensing, as they are expressed in pheromone-sensitive olfactory sensory neurons. There are three SNMP genes reported from Lepidoptera (28), with SNMP1 and SNMP2 best characterized. SNMP1 expression is reported as being limited to moth antennae (*Antheraea polyphemus* (Rogers et al., 1997); *M. sexta* (Rogers et al., 2001), but SNMP1 orthologs have more recently been reported from other chemosensory appendages in insects (29). Contrary to the reported restriction of SNMP1 to the antenna, we detected a putative ortholog, *g9123*, from at least one sample of all tissues except the male proboscis. However, there is a large amount of variation in the number of reads detected across samples, suggesting this finding should be validated with PCR. SNMP2 expression is more widespread, being detectable in multiple body parts throughout development in *B. mori* (28), but shows highest expression in antennal tissue of moths (29,75). Here, an SNMP2 ortholog, *g4539*, showed significantly greater expression in the *H. lineata* ovipositor when compared with male genitalia. To date, there is no evidence that SNMPs play a role in sensing host plant compounds. But given that plant surfaces are covered in long-chain fatty acids and SNMPs putatively function in long-chain fatty acid sensing, our data suggest this should be studied further.

### OBPs as candidate oviposition-relevant genes

Like SNMPs, OBPs were first characterized by their role in pheromone sensing (56,76) but have since been implicated in multiple chemosensory contexts (27,76–78). They are small (100 - 160 amino acids), water-soluble proteins that are secreted by accessory cells around olfactory receptor neurons and exist in high concentrations in the sensilla hemolymph (79). OBPs solubilize and transport hydrophobic ligands through the aqueous hemolymph to ORs, influencing the sensitivity of odor transduction (80). Insect OBPs vary greatly in sequence homology (22) with some showing less than 20% sequence similarity across genera (81), suggesting they may play a role in odorant discrimination through selective binding (80).

Odorant binding proteins have been implicated in oviposition behavior insects through their role in detecting host plant compounds. An ortholog of *DmelOBP99a* has been implicated in oviposition behavior in the oriental fruit fly, *Bactrocera dorsalis*. Specifically, silencing this gene reduced egg-laying on a preferred host while increasing egg-laying on a non-preferred host (82). In the swallowtail butterfly *Atrophaneura alcinous*, an OBP with female-specific expression in the legs acts as an oviposition stimulant, potentially through binding specific compounds the butterfly uses to detect preferred host plants (83). In our data, neither of the OBPs that showed higher expression in the ovipositor and female legs have orthologs definitively associated with oviposition behavior in other insects. One of them, *g9036,* clusters closely with *OBP19d* orthologs, an abundant, widely expressed OBP in *D. melanogaster* (58). The second one, *g9040*, is most closely related to the OBP *LUSH*, which is implicated in pheromone-mediated behaviors in *D. melanogaster* (56), but likely also plays a role in detecting other odorants (57,84). This differentially expressed OBP could be involved in mating-related behaviors, rather than or in addition to oviposition.

## Conclusions

The aim of our study was to identify genes in *H. lineata* that might be involved in host plant associations. Here, we provide the first annotation of chemosensory genes in *H. lineata* and suggest several candidates for future studies on sex-specific chemosensory behaviors. As with other differential gene expression studies, our data serve as an important piece of evidence in a much broader, growing literature on this species. Because variables such as the age of moths and time of year they are collected might influence DGE results, confidence in these findings will be enhanced by repeated experiments with a greater number of replicates.

## Additional Files

Godfrey_et_al_figures_supplemental.docx contains supplemental figures cited within the manuscript.

Godfrey_et_al_tables_supplemental.xlsx contains supplemental tables with specimen, sample, and chemosensory gene annotation information cited within the manuscript.

## Abbreviations

DEG: differentially expressed genes
DE: differentially expressed
DGE: differential gene expression
GO: gene ontology
GR: gustatory receptor
IR: ionotropic receptor
OG: orthologous group
OR: odorant receptor
OBP: odorant binding protein
SNMP: sensory neuron membrane proteins
PCR: polymerase chain reaction
TPM: Transcripts per Million

## Declarations

### Ethics approval and consent to participate

While the project described here did not involve humans or other vertebrates, we acknowledge that insects exhibit stress upon being handled and that data suggest insect populations are threatened regions of the world. We therefore collected as few specimens as possible for our studies, retained them in a cool, dark place after capture and during transport, and euthanized them quickly by freezing.

### Availability of data and materials

RNA-seq raw reads used for this paper are available in the NCBI Sequence Read Archive under bioproject PRJNA1145426. The analysis pipeline for differential gene expression and all files used for functional annotation are available from GitHub: https://github.com/rkeatinggodfrey/Hyles_lineata_chemosensory_gene_expression

### Author Contributions

RKG and AYK conceptualized the project, RKG, PD, and AA developed the methodology, RKG, EC, RH, KK, CM, FM, and GP conducted formal analyses of the data, RKG wrote the original draft and all other authors reviewed it and provided critical edits, RKG and GP contributed visualizations, RKG, AA, and PD supervised the project, RKG conducted project administration, RKG and AYK performed funding acquisition.

## Acknowledgements

We are thankful to Goggy Davidowitz and Sarah Britton for their knowledge and enthusiasm for *Hyles lineata* and their help collecting moths. We are also grateful to the U.S. Department of Agriculture’s Forest Service for providing temporary use permit for the Coronado National Forest to collect moths for this project.

Initial data analysis for this work was performed by 31 undergraduate students in a bioinformatics and sensory biology course-based undergraduate research experience (CURE) at the University of Florida. We are very grateful to the all of those students for their contribution to this project; those who are not co-authors on this publication are: Mak Aganovic, Hannah Amaya, Yoan Argote, Ryan Banner, David Castellanos, Daniel Cho, Omie Coyne, Olivia Drake, Peter Emery, Audrey-Anne Haynes, Aiden Hussey, Isabella Koushiar, Lauren Lambie, Maryanne Moore, Lauryn Murphy, Kailie O’Neal, Dhruv Panwar, Alondra Peña, Alyson Port, Nicole Preil, Ryan Rodriguez, Margaret Shealy, Joshua Siu, Katherine Timmins, and Myan Tran. For support of this CURE course, we thank the University of Florida Entomology and Nematology Department, the UF College of Agricultural and Life Sciences, the UF Center for Undergraduate Research, and the UF International Center Office of Global Learning.

