## Supplemental figures for "Sex-specific chemosensory gene expression in the whitelined sphinx moth (Lepidoptera: *Hyles lineata*) suggests a role for odorant-binding proteins in host plant choice"

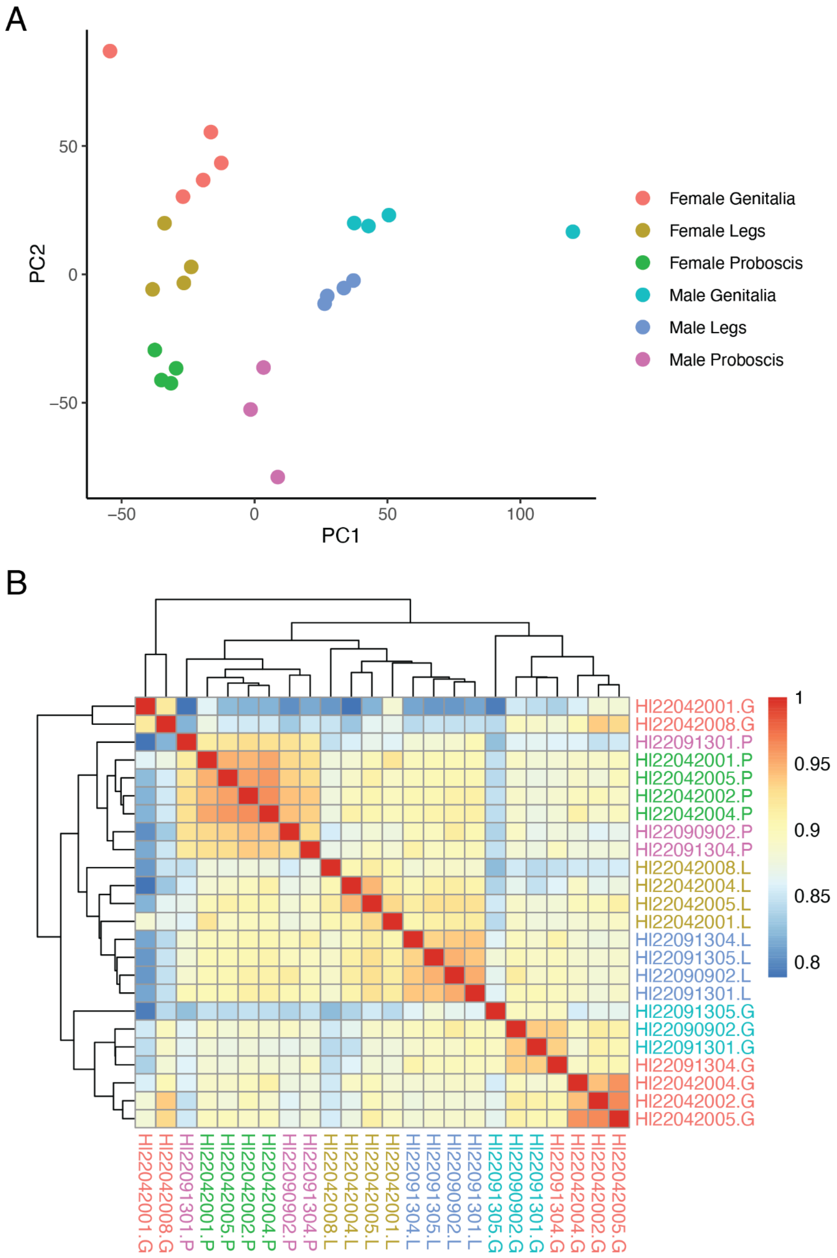


Figure S1. (A) Principal components analysis and (B) hierarchical clustering heat map of pairwise correlations for samples used for differential gene expression analysis. Correlation values indicated by the continuous color scale shown on the right. Color of sample names in (B) coincide with those indicating sex and body part in (A).

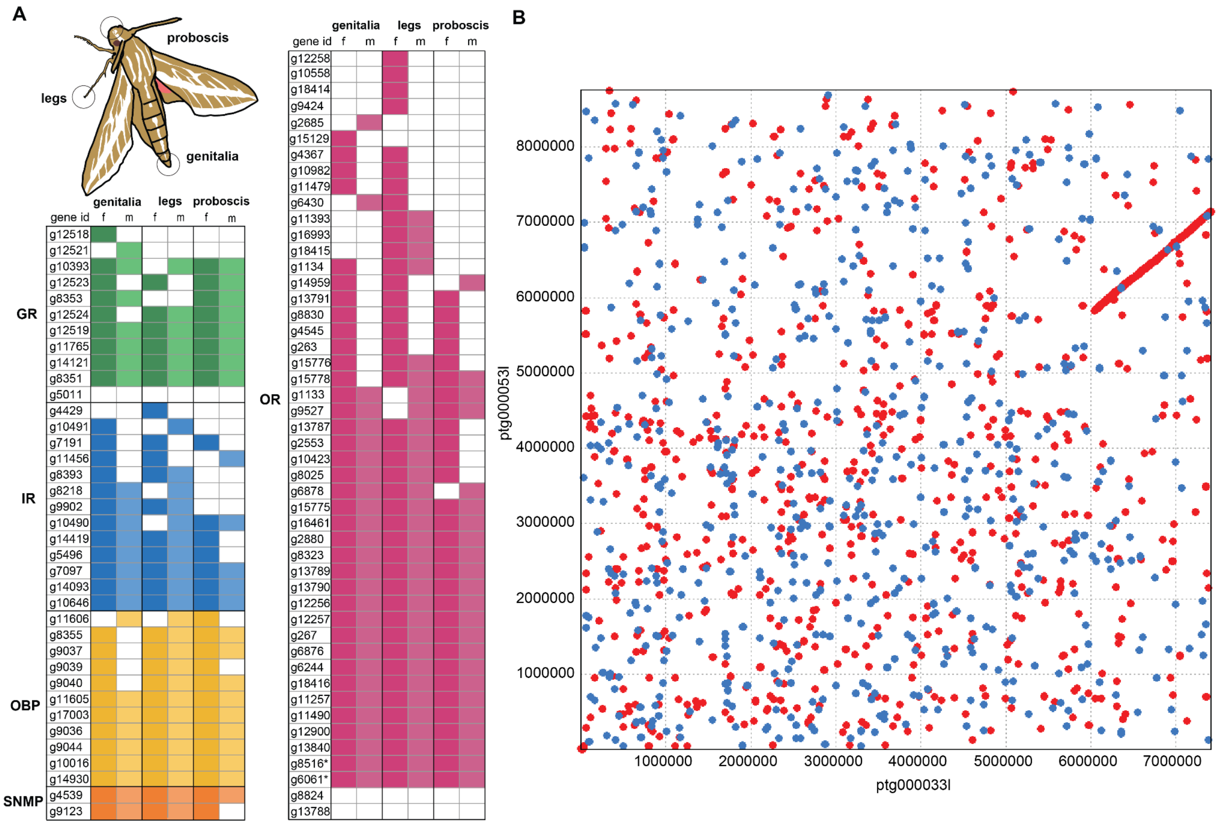


Figure S2. (A) Annotation of chemosensory genes. Presence (filled cells) or absence (empty cells) of annotated genes in gene expression data. Rows show gene IDs from a structural annotation. Filled cells indicate that the gene was detected in ≥ one sample of each type (body part and sex). Genes with only blank cells were functionally annotated, but not recovered from any samples. Asterisks indicate the two orco genes detected in annotation. (B) Synteny analysis showing two H. lineata assembly contigs () that each contain an Orco sequence. There is an area of approximately 1250 kb showing sequence alignment that indicates an uncollapsed region in the assembly.

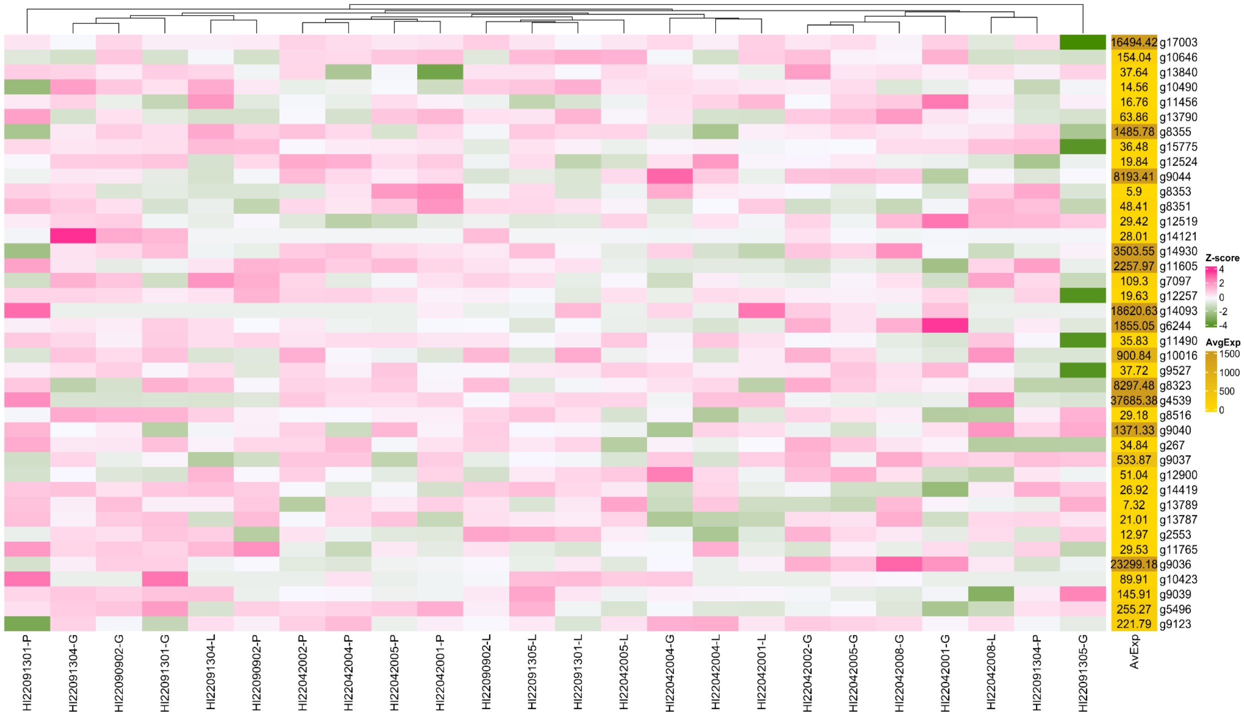


Figure S3. Chemosensory gene expression in *Hyles lineata* samples assessed with differential gene expression using DESeq2. Samples IDs appear across the bottom and gene IDs to the right.
